# Self-navigating the “Island of Reil”: a systematic review of real-time fMRI neurofeedback training of insula activity

**DOI:** 10.1101/2022.03.07.483236

**Authors:** Yuan Zhang, Qiong Zhang, Benjamin Becker, Keith M. Kendrick, Shuxia Yao

## Abstract

Real-time fMRI (rtfMRI) neurofeedback (NF) is a novel noninvasive technique that permits individuals to voluntarily control brain activity or connectivity, with demonstrated feasibility in experimental and therapeutic applications. The crucial role of the insula in emotional and salience processing makes it a popular target in rtfMRI studies although there is no systematic review of their efficacy. To promote our understanding of mechanisms underlying insula regulation and thereby facilitate therapeutic translation, the present review identified 22 preclinical and clinical studies and found that rtfMRI-based NF training is efficient for modulating insula activity (> 86%) and its associated behavioral and neural changes. Based on findings, continuous feedback for multiple training sessions, specific strategies validated before application, inclusion of a practice session, and choosing appropriate localization strategies are suggested as guidelines. We also recommend standardization of training protocol design, statistical analyses and results reports for future studies. In summary, the present review informs both fundamental research and therapeutic translation of NF training as an intervention in mental disorders, particularly those with insula dysfunction.

## 1. Introduction

The insula, known as the “Island of Reil”, is located deep in the lateral sulcus (Flynn et al., 1999) and extensively interconnected with other brain regions (Flynn et al., 1999; Saper, 2002). In line with this extensive interconnection, the insula represents an integrative hub in the emotion and salience network (Fox et al., 2005; Menon and Uddin, 2010) and plays a vital role in the integration of external-oriented sensory input and internal-oriented awareness (Craig, 2009, 2002). The insula is a cytoarchitectonically and functionally heterogenous region organized along a posterior-to-anterior gradient such that the anterior insula (AI) is mainly involved in interoceptive processing (Caseras et al., 2013; Critchley, 2004; Simmons et al., 2013), emotion (Lamm and Singer, 2010), risky decision-making (Clark et al., 2008) and empathy (Fan et al., 2011; Valentini, 2010), while the posterior insula (PI) primarily integrates somatosensory, vestibular, and motor information by receiving afferent projections from the spinal cord and brainstem via the thalamus (Flynn et al., 1999). Functional and structural dysregulations of the insular cortex are closely associated with various mental disorders, including anxiety disorders (Klumpp et al., 2012; Shah et al., 2009), major depression disorders (MDD) (Wiebking et al., 2015, 2010; Xu et al., 2020), autism spectrum disorders (Ebisch et al., 2011; Francis et al., 2019; Uddin and Menon, 2009), schizophrenia (Caldiroli et al., 2017; Tuominen et al., 2017; Wylie and Tregellas, 2010), addiction (Droutman et al., 2015; Klugah-Brown et al., 2020; Naqvi and Bechara, 2009), alexithymia (Ernst et al., 2013; J. Li et al., 2019) and Parkinson’s disease (Christopher et al., 2014; Criaud et al., 2016). Thus, a noninvasive brain modulation approach that can directly target the insula and normalize its dysregulations may represent a novel and transdiagnostically promising therapeutic strategy.

Real-time fMRI (rtfMRI) neurofeedback (NF) training capitalizes on recent progress in fMRI techniques and represents a noninvasive approach to modulate regional activity or functional connectivity in cortical and subcortical brain regions (Paret et al., 2019; Watanabe et al., 2017; Weiskopf, 2012; Zhao et al., 2019). More specifically, integrated within a training rtfMRI NF can assist participants to gain voluntarily control over brain activity by transforming the BOLD signal into visual real-time feedback to guide the acquisition of neural self-regulation. Previous studies have demonstrated that this approach allows individuals to gain self-regulation over several brain regions, including the amygdala (Barreiros et al., 2019; Young et al., 2017b), the anterior cingulate cortex (ACC) (Hamilton et al., 2011; Zilverstand et al., 2017), the prefrontal cortex (Kohl et al., 2019; Mayeli et al., 2020; Sherwood et al., 2016), the hippocampus (Zhu et al., 2019), as well as the insula (Caria et al., 2010, 2007). The clinical application potential of rtfMRI NF training has also been supported by accumulating studies in clinical populations, including MDD (Young et al., 2014; Zotev et al., 2020), anxiety disorder (Scheinost et al., 2013; Zilverstand et al., 2015), borderline personality disorder (Paret et al., 2016; Zaehringer et al., 2019), addiction (Canterberry et al., 2013; Kirschner et al., 2018), post-traumatic stress disorder (PTSD) (Nicholson et al., 2017; Zweerings et al., 2018), and attention deficit hyperactivity disorder (Alegria et al., 2017; Zilverstand et al., 2017). Based on the promising potential in both, basic research and therapeutic application, the rtfMRI NF field has progressed rapidly during the last decade. However, despite the promising translational potential of rtfMRI most therapeutic applications remain in the experimental stage. Thus, a systematic review of key characteristics of NF training and its associated behavioral and neural changes is important for facilitating translation of rtfMRI into therapeutic application.

Given the crucial role of the insula in emotional and salience processing (Lamm and Singer, 2010; Menon and Uddin, 2010), it has been one of the most popular regions that have been investigated in the field of rtfMRI both in healthy (e.g., Caria et al., 2010, 2007; Yao et al., 2016) and clinical populations including schizophrenia (Ruiz et al., 2013), alcohol use disorders (Karch et al., 2015), MDD (Linden et al., 2012), and Parkinson’s disease (Tinaz et al., 2018). Surprisingly, to date no review has systematically summarized progress in the field of rtfMRI-based insula modulation. Previous reviews primarily focus on general progress in the field of rtfMRI (e.g., Paret et al., 2019; Thibault et al., 2018; Weiskopf, 2012), the general therapeutic potential of rtfMRI NF training in mental disorders (e.g., Dudek and Dodell-Feder, 2021; Tursic et al., 2020) or in specific disorders such as addiction (Martz et al., 2020), schizophrenia (Gandara et al., 2020) and eating disorders (Bartholdy et al., 2013), except two reviews specifically focusing on progress of rtfMRI NF training on the amygdala during emotion regulation (Barreiros et al., 2019) and in MDD (Young et al., 2018).

Against this background, the present review conducted a systematic review of the efficacy of rtfMRI NF training targeting the insula, its associated behavioral and neural effects, and factors and parameters that may affect training success. More specifically, the main objectives of the present review are providing a systematic overview on: (1) What characteristics (e.g., training protocol design, feedback types, regulation strategies, etc.) were used in NF training protocols of rtfMRI NF training studies on insula? (2) Can rtfMRI NF training lead to consistently successful volitional control of insula, and if so, can this training success be maintained to a “transfer” session without NF? and (3) What behavioral and neural changes can be induced by rtfMRI NF training of the insula? It has to be noted that given the publication bias that most of the available studies were based on statistically significant findings, ratio of training success is thus not an indicator of training efficacy in the whole population of studies (see also Barreiros et al., 2019). Consequently, comparisons between specific characteristics that may affect training success of insula regulation are not informative. Therefore, in the present study we did not conduct statistical analyses comparing key characteristics in the training protocol.

## 2. Methods

### 2.1. Literature Search and Inclusion Criteria

Literature search was conducted by two authors independently in line with the PRISMA guidelines (Liberati et al., 2009; Moher et al., 2009) (Fig. 1). Specifically, literature on rtfMRI NF training on the insula was identified using the Web of Science and PubMed databases with the search terms “neurofeedback” and “fMRI” or “functional magnetic resonance imaging” or “functional MRI” and “insula” or “insular”. A second search was then performed in these two databases with the following keywords: “rtfMRI” or “real time” or “real-time” and “fMRI” or “functional magnetic resonance imaging” or “functional MRI” and “insula” or “insular”. Literature search was undertaken in March 2021 and resulted in 1284 articles (902 articles in PubMed and 382 articles in Web of Science).

**Fig. 1.**
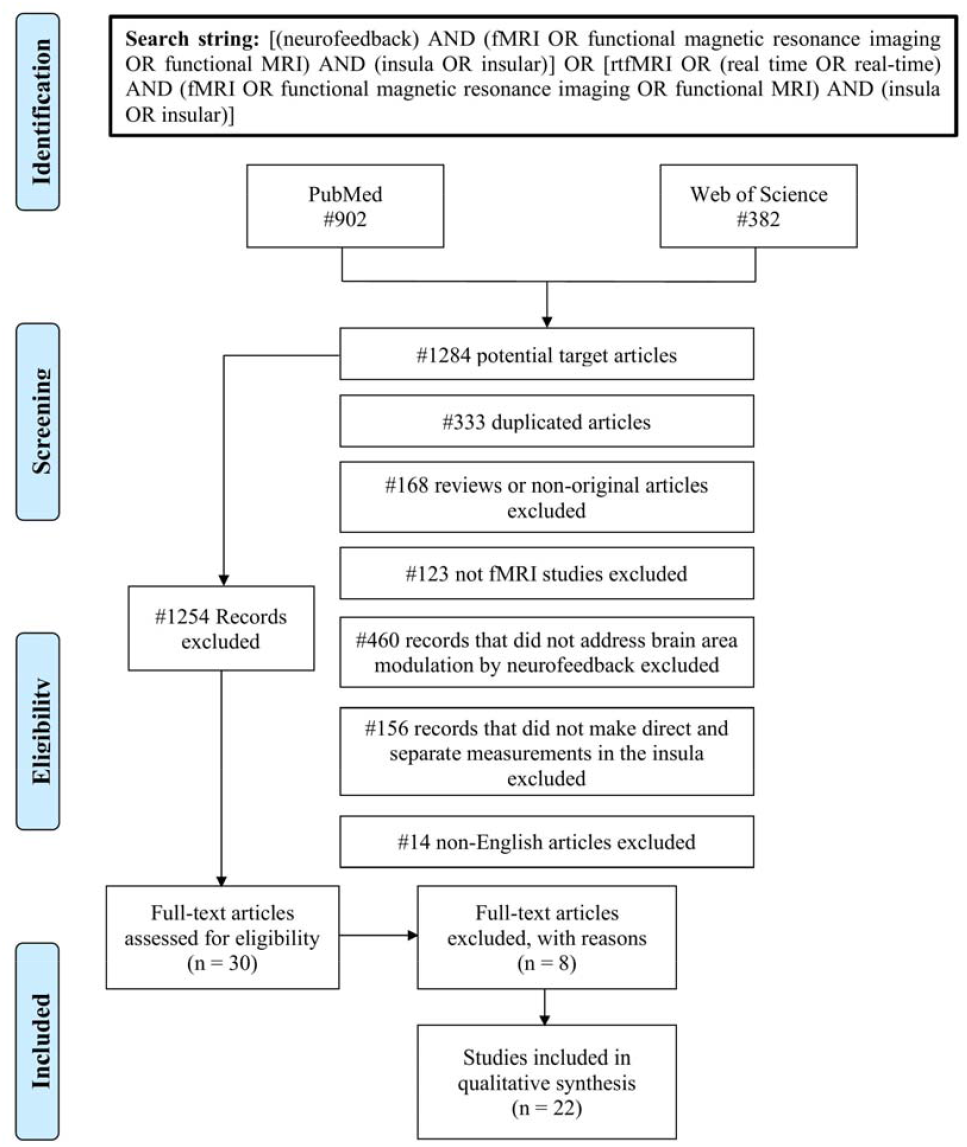
Flowchart of the literature selection procedure based on the PRISMA guidelines.

The 1284 articles were further screened by analyzing titles and abstracts based on the following study eligibility criteria: (1) written in English; (2) utilized rtfMRI-NF training (rather than other neurofeedback modalities such as electroencephalography, functional near-infrared spectroscopy and electrophysiology); (3) used BOLD signal of the insula (or its connectivity) as neurofeedback sources; (4) reported original research (i.e. reviews or commentaries were excluded). Based on these criteria, 1254 articles were excluded, including 333 duplicated articles, 14 non-English articles, 123 non-fMRI articles, 460 non-neurofeedback fMRI articles, 156 rtfMRI articles using other brain regions as neurofeedback sources, and 168 non-original research articles. The remaining 30 articles were entered into the next stage for full text analyses, in which another 8 articles were excluded due to the training not being based on the BOLD signal of the insula (Li et al., 2016b; Ninaus et al., 2013; Weaver et al., 2020; Zhang et al., 2020), no quantitative results being reported (Emmert et al., 2017a), focus on evaluation of task-related BOLD signal artifacts during rtfMRI NF training (Zhang et al., 2011), or conducting additional analyses based on data from pervious rtfMRI studies (Caria, 2020; Lee et al., 2011). These steps resulted in 22 articles for the final quantitative analysis.

### 2.2. Data Extraction

The following characteristics were extracted from each of the 22 included articles: (1) demographic information of participants: population (healthy/clinical), sample size, age and gender distribution; (2) feedback characteristics: feedback source (single ROI/functional connectivity of the insula), feedback types (intermittent/continuous feedback); (3) regulation strategies and up/down regulation; (4) localization methods of target regions: anatomical/functional localization (in case of functional localizer the task paradigm); (5) protocol design: number of training sessions, number and duration of regulation blocks for each session, practice and transfer session (practice and transfer conducted or not), control group (whether a control group was included or not), and stimuli (whether other stimuli were displayed in addition to the feedback display); (6) outcome measures and main findings (training effects on activity or functional connectivity of target regions and associated neural and behavioral changes). Training success on self-regulation of the insula activity and associated neural and behavioral effects were determined by comparing differences between the experimental and control groups, between regulation and rest blocks, or between the first and last sessions. Other training-related measurements, such as resting state functional connectivity (rsFC), task-based performance and clinical symptoms, were examined by comparing differences between pre- and post-training.

## 3. Results

### 3.1. Participants

The final 22 qualified articles consisted of 13 studies in healthy and 9 in clinical populations (**Table 1** and **Fig. 2A**). Clinical studies included participants with schizophrenia (Ruiz et al., 2013), obsessive-compulsive disorder (OCD) (Buyukturkoglu et al., 2015), spider phobia (Zilverstand et al., 2015), MDD (Linden et al., 2012), alcohol and tobacco addiction (Karch et al., 2019, 2015; Rana et al., 2020), Parkinson’s disease (Tinaz et al., 2018), and criminal psychopaths (Sitaram et al., 2014). Significant training effects on insula self-regulation were found in 12 of 13 studies (92.31%) in healthy subjects and 7 of 9 studies in clinical populations (77.78%) (**Fig. 3**). Of note the 2 clinical studies showing no significant training effects were conducted in comparably small samples and in patient groups characterized by marked regulatory control deficits, i.e. OCD patients (n = 3; Buyukturkoglu et al., 2015) and criminal psychopaths (n = 4; Sitaram et al., 2014).

**Fig. 2.**
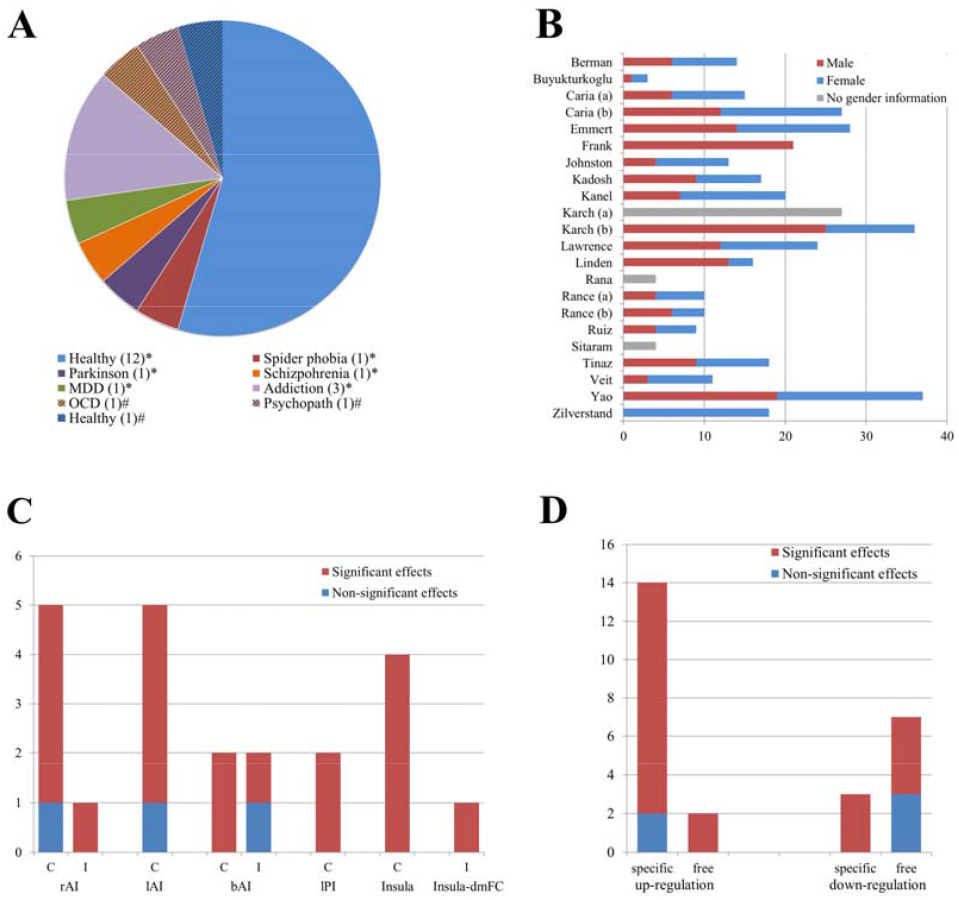
Population (A) and gender distribution (B) in the qualified 22 studies. (C) Feedback types categorized by target regions. (D) Types of regulation strategies categorized by regulation directions. Note. Abbreviations in alphabetical order: AI = Anterior insula; b = bilateral; C = Continuous feedback; I = Intermittent feedback; Insula-dmFC: Functional connectivity between the insula and dorsomedial frontal cortex; l = left; PI = Posterior insula; r = right; * = Studies showing significant effects on insula regulation; # = Non-significant training effects on insula regulation.

**Fig. 3.**
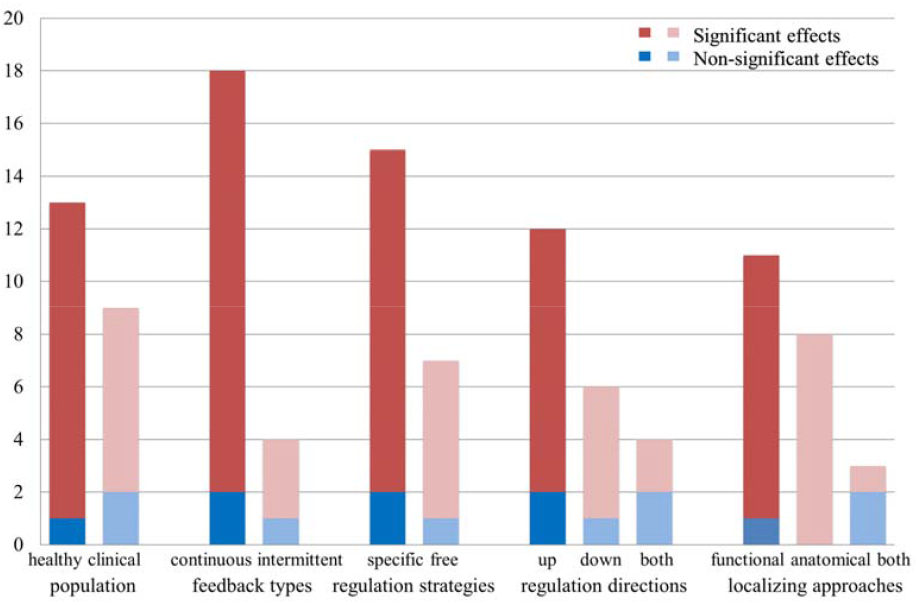
Proportion of studies showing significant training effects on insula regulation according to population, feedback types, regulation strategies, regulation directions and localizing approaches.

**Table 1.** Participants’ demographics, feedback and regulation strategies.

| Study | Population | N (F) | Age M (SD) | Target ROIs | Feedback types | Regulation direction | Regulation strategies |
| --- | --- | --- | --- | --- | --- | --- | --- |
| Berman et.al (2013) | Healthy | 14 (8) | 29.7 (8.2) | rAI | Continuous | Up | Mental imagery |
| Buyukturkoglu et.al (2015) | OCD | 3 (2) | 23.33 (3.09) | bAI | Intermittent | Down | Free |
| Caria et.al (2007) | Healthy | 15 (9) | 25.13 (-) | rAI | Continuous | Up | Mental imagery |
| Caria et.al (2010) | Healthy | 27 (15) | 27.51 (-) | lAI | Continuous | Up | Mental imagery |
| Emmert et.al (2014) | Healthy | 28 (14) | 27.5 (2.3) | lAI# | Continuous | Down | Free |
| Frank et.al (2012) | Healthy(obese/lean) | 21 (0) | 26.24 (0.86) | bAI | Continuous | Up | Mental imagery |
| Johnston et.al (2010) | Healthy | 13 (9) | 21-52 | Insula/amygdala/VLPFC | Continuous | Up | Mental imagery |
| Kadosh et.al (2016) | Healthy | 17 (8) | 11.6 (2.5) | rAI | Continuous | Up/Down | Mental imagery |
| Kanel et.al (2019) | Healthy | 20 (13) | 24.6 (3.32) | rAI | Continuous | Up | Emotional voice empathizing |
| Karch et al (2015) | Alcohol addiction and healthy | 27 (-) | 18-60 | Insula/ACC/DLPFC | Continuous | Down | Free |
| Karch et al (2019) | Tobacco addiction(abstinence/relapse) | 36 (11) | 43.83 (12.37) | Insula/ACC/DLPFC | Continuous | Down | Free |
| Lawrence et.al (2014) | Healthy | 24 (12) | 27.33 (4.71) | rAI | Continuous | Up | Mental imagery/Body awareness |
| Linden et.al (2012) | MDD | 16 (3) | 48.44 (12.99) | Insula/DLPFC | Continuous | Up | Mental imagery |
| Rana et.al (2020) | Nicotine addiction | 4 (-) | - | bAI | Intermittent | Down | Free |
| Rance et.al (2014a) | Healthy | 10 (6) | 27.8 (4.71) | lPI/rACC | Continuous | Up/Down | Free |
| Rance et.al (2014b) | Healthy | 10 (4) | 29 (6.48) | lPI# | Continuous | Up/Down | Free |
| Ruiz et.al (2013) | Schizophrenia | 9 (5) | 26.3 (4.5) | bAI | Continuous | Up | Mental imagery |
| Sitaram et.al (2014) | Criminal with psychopaths | 4 (-) | 31.5 (3.5) | lAI | Continuous | Up | Mental imagery |
| Tinaz et.al (2018) | Parkinson’s disease | 18 (9) | 64.49 (9.74) | Insula-dmFC | Intermittent | Up | Mental imagery |
| Veit et.al (2012) | Healthy | 11 (8) | 21-28 | lAI | Continuous | Up/Down | Mental imagery |
| Yao et.al (2016) | Healthy | 37 (18) | 21.97 (1.4) | lAI | Continuous | Up | Mental imagery |
| Zilverstand et.al (2015) | Spider phobia | 18 (18) | 21.2 (1.78) | rAI | Intermittent | Down | Cognitive reappraisal |

*Note.* Abbreviations in alphabetical order: ACC = Anterior cingulate cortex; AI = Anterior insula; b = Bilateral; DLPFC = Dorsolateral prefrontal cortex; F = Female’s number; l = Left; M = Mean; MDD = Major depressive disorder; N = Numbers of subjects; OCD = Obsessive compulsive disorder; PI = Posterior insula; r = Right; SD = Standard deviation; VLPFC = Ventrolateral prefrontal cortex; - = No reported results in articles; # = Two training groups with different target ROIs.

There were 382 participants in total in these qualified studies and sample size ranged from 3-37, with 4 clinical studies encompassing less than 10, 10 studies with a range of 10 to 19, 6 studies from 20 to 30, and 2 studies more than 30 (**Fig. 2B**). For 19 studies information on gender distribution was available and the percentage of females was 49.57% with 17 studies including both male and female participants while two studies only recruited male (Frank et al., 2012) or female participants (Zilverstand et al., 2015). The remaining 3 studies did not report gender information (Karch et al., 2015; Rana et al., 2020; Sitaram et al., 2014). For age, 3 studies only reported age range and one study did not report specific age information (Johnston et al., 2010; Karch et al., 2015; Rana et al., 2020; Veit et al., 2012). The mean age of the remaining 18 studies is 30.31 years old. While the youngest sample (mean = 11.6) was observed in a study examining the rtfMRI training effects on the emotion regulation network in adolescents (Cohen Kadosh et al., 2016), the most senior sample (mean = 64.49) was found in a clinical study on patients with Parkinson’s disease (Tinaz et al., 2018). All details of participants’ characteristics are presented in **Table 1** and **Fig. 2**.

### 3.2. Feedback types

RtfMRI neurofeedback is normally displayed using a visual feedback signal (e.g., in the form of a thermometer bar or using magnitude of momentary reward as feedback of performance) intermittently or continuously reflecting regional activity or functional connectivity strengths (e.g., Berman et al., 2013; Rana et al., 2020; Yao et al., 2016). Apart from one study employing functional connectivity-based NF between the insula and the dorsomedial frontal cortex (dmFC) (Tinaz et al., 2018), the other 21 studies displayed feedback based on regional specific BOLD signal amplitude of the insula, which included 6 studies focusing on the right AI, 5 studies focusing on the left AI, 4 studies focusing on bilateral AI, 2 studies focusing the left PI, and 4 studies focusing on the whole insula (**Table 1** and **Fig. 2C**). With respect to the feedback type, 18 studies employed continuous feedback, which was continuously presented to participants in regulation/rest blocks. The remaining 4 studies used intermittent feedback presented at the end of each regulation/rest block (**Table 1**). Significant training effects were found in 3 of 4 studies (75%) using intermittent feedback and 16 of 18 studies (88.89%) using continuous feedback (**Fig. 3**).

### 3.3. Regulation strategies and direction

Efficient NF training commonly depends on valid regulation strategies which can be specific following instruction or freely selected by participants. Among the 22 qualified studies, participants in 7 studies were instructed to find their own strategy (i.e. free strategies) to effectively regulate insula activation (Buyukturkoglu et al., 2015; Emmert et al., 2014; Karch et al., 2019, 2015; Rana et al., 2020; Rance et al., 2014a, 2014b). Fifteen studies provided specific regulation strategies with mental imagery being the most popular strategy (13 studies), including autobiographical memory recall (Berman et al., 2013; Caria et al., 2007; Lawrence et al., 2014; Linden et al., 2012; Ruiz et al., 2013; Sitaram et al., 2014; Yao et al., 2016), emotional recall (Cohen Kadosh et al., 2016; Frank et al., 2012; Veit et al., 2012), and motor imagery (Tinaz et al., 2018). The remaining two studies used emotional voice empathizing (Kanel et al., 2019) or cognitive reappraisal (Zilverstand et al., 2015) as their regulation strategies. Significant training effects were observed in 6 of 7 studies (85.71%) using free strategies and in 13 of 15 studies (86.67%) using specific strategies (**Fig. 3**). In contrast to regulation blocks, to induce brain activation back to baseline level, participants were asked to close eyes and rest, to relax, focus on a cross, count number or view neutral stimulus during rest blocks.

With respect to regulation direction, 12 studies aimed at up-regulation (including one study on increasing the insula-dmFC functional connectivity), 6 studies on down-regulation, and 4 studies on both up- and down-regulation of insula activity (**Fig. 3**). Up- and down-regulation studies had a comparable training success rate (83.33%), with 10 of the 12 up-regulation studies (Caria et al., 2010, 2007; Frank et al., 2012; Johnston et al., 2010; Kanel et al., 2019; Lawrence et al., 2014; Linden et al., 2012; Ruiz et al., 2013; Tinaz et al., 2018; Yao et al., 2016) and 5 of 6 down-regulation studies reporting successful regulation of insula activity (Emmert et al., 2014; Karch et al., 2019, 2015; Rana et al., 2020; Zilverstand et al., 2015). For the 4 studies involving both up- and down-regulation, all of them reported successful up-regulation (Cohen Kadosh et al., 2016; Rance et al., 2014a, 2014b; Veit et al., 2012) and half of them reported successful down-regulation (Rance et al., 2014a, 2014b) of the insula activity (**Fig. 2D**). Thus, in total there were 14 studies reporting training success (87.50%) among the 16 studies involving up-regulation and 7 studies reporting training success (70%) among the 10 studies involving down-regulation (**Fig. 3**). In contrast to 2 in 16 up-regulation studies applying free regulation strategies in insula modulation, most of the down-regulation studies used free regulation strategies (7 in 10 studies) (details see **Table 1**). While there were 9 studies in the 10 down-regulation studies presenting stimuli to participants during regulation blocks, only 5 in 16 up-regulation studies did the same (details see **Table 2**).

**Table 2.** Characteristics of neurofeedback training protocol design.

| Study | Feedback in control groups | Practice session | Transfer session | Training sessions |  | Stimuli (Ratings) | Time point of stimuli delivery | Localizer | Localizer tasks |
| --- | --- | --- | --- | --- | --- | --- | --- | --- | --- |
|  |  |  |  | Sessions | Blocks (Duration) |  |  |  |  |
| Berman et al. (2013) | / | Yes | Yes | 4 | 5 (30) | / | / | both | Suppression task vs. rest |
| Buyukturkoglu et al. (2015) | / | / | Yes | 16# | 6 (27) | Disgusting pictures (Yes) | Loc/Training\$/Pre-Post | both | Disgust vs. neutral pictures |
| Caria et al. (2007) | Sham/No feedback | / | Yes | 4 | 4 (22.5) | / | / | anatomical | / |
| Caria et al. (2010) | Sham/No feedback | / | / | 4 | 5 (30) | Aversive pictures (Yes) | Training\$ | anatomical | / |
| Emmert et al. (2014) | / | / | / | 4 | 4 (30) | Heat pain (Yes) | Loc/Training\$ | functional | Pain stimulation vs. rest |
| Frank et al. (2012) | / | / | / | 6# | 7 (30) | / | / | anatomical | / |
| Johnston et al. (2010) | / | / | / | 3 | 12 (20) | Emotional pictures (No) | Loc | functional | Negative vs. neutral pictures |
| Kadosh et al. (2016) | / | / | / | 4 | 9 (20) | Face pictures (No) | Loc | functional | Emotional go/no-go |
| Kanel et al. (2019) | Sham feedback | / | Yes | 16# | 8 (24) | Emotional voice (No) | Training\$ | anatomical | / |
| Karch et al. (2015) | Sham feedback | / | / | 4 | 4 (40) | Alcohol pictures (No) | Loc/Training\$ | functional | Alcoholic vs. neutral pictures |
| Karch et al. (2019) | Sham feedback | / | / | 9# | 4 (40) | Tobacco pictures (No) | Loc/Training\$ | functional | Tobacco vs. neutral pictures |
| Lawrence et al. (2014) | Sham feedback | / | / | 4 | 5 (30) | Affective pictures (Yes) | Pre-Post | anatomical | / |
| Linden et al. (2012) | No feedback | / | / | 12# | 12 (20) | Emotional pictures (No) | Loc | functional | Emotional pictures vs. rest |
| Rana et al. (2020) | / | / | Yes | 8# | 6 (26) | Nicotine pictures (No) | Training\$/Pre-Post | anatomical | / |
| Rance et al. (2014 a) | / | Yes | / | 24# | 6 (45) | Heat pain (Yes) | Training\$ | anatomical | / |
| Rance et al. (2014 b) | / | Yes | / | 24# | 6 (45) | Heat pain (Yes) | Training\$ | anatomical | / |
| Ruiz et al. (2013) | / | / | Yes | 12# | 6 (30) | Face pictures (Yes) | Post | functional | Emotional recall vs. rest |
| Sitaram et al. (2014) | / | / | / | 12# | 6 (30) | Aversive pictures (Yes) | Pre-Post | functional | Emotional recall vs. count number |
| Tinaz et al. (2018) | No feedback | Yes | Yes* | 10-12# | 5 (40) | / | / | both | Heartbeat detection vs. rest |
| Veit et al. (2012) | / | / | / | 3 | 12 (9) | Aversive pictures (No) | Loc/Training\$ | functional | Aversive vs. neutral pictures |
| Yao et al. (2016) | Sham feedback | / | Yes* | 4 | 5 (30) | Painful pictures (Yes) | Loc/Training | functional | Painful vs. non-painful pictures |
| Zilverstand et al. (2015) | No feedback | Yes | / | 3 | 4 (12.5) | Spider pictures (Yes) | Loc/Training\$ | functional | Spider pictures vs. rest |

*Note.* Abbreviations in alphabetical order: Loc = Stimuli were used in functional localization; Pre-Post = Stimuli were presented pre-training and post-training; # = Training sessions were performed across different days; \$ = Stimuli were presented during regulation blocks.

### 3.4. Localizing approaches of target regions

Accurate localization of the target region is vital for precise feedback and the corresponding regulation strategies. Approaches for target region localization can be categorized into anatomical and functional localization. In the present study, there were 8 studies employing anatomical localization (Caria et al., 2010, 2007; Frank et al., 2012; Kanel et al., 2019; Lawrence et al., 2014; Rana et al., 2020; Rance et al., 2014a, 2014b), 11 studies applying functional localization (Cohen Kadosh et al., 2016; Emmert et al., 2014; Johnston et al., 2010; Karch et al., 2019, 2015; Linden et al., 2012; Ruiz et al., 2013; Sitaram et al., 2014; Veit et al., 2012; Yao et al., 2016; Zilverstand et al., 2015) and 3 studies combining both approaches to define the insula (Berman et al., 2013; Buyukturkoglu et al., 2015; Tinaz et al., 2018). We categorized the different tasks used for functional localization into 3 main types, including emotional recall tasks, passive observation task (emotional/addictive pictures) and other specific tasks (e.g., blink suppression, emotional go/no-go, and heartbeat detection and so on) (see Table 2). While all of the 8 studies using anatomical localization showed significant training effects on insula activity, 11 of the 14 studies (78.57%) applying functional localization reported significant training effects on the insula regulation (**Fig. 3**). Note that 2 of the 3 studies showing no significant training effects were the same to the 2 studies with low sample sizes in OCD patients (n = 3) and criminal psychopaths (n = 4) as discussed in the “Participants” section (Buyukturkoglu et al., 2015; Sitaram et al., 2014).

### 3.5. Protocol design

#### 3.5.1. Training sessions

Training protocols in the 22 qualified studies employed between 3 to 24 sessions (mean = 8.73), with half of the studies incorporating 3 or 4 training sessions performed on a single day and the other half (including 7 clinical studies) comprising 6 or more sessions and being performed from 2 to 4 separate days (**Table 2**). Sessions included in clinical studies were slightly more than in healthy subject studies (9.78 vs. 8 sessions respectively). Each session included 4 to 12 regulation blocks (mean = 6.41) alternating with rest blocks. The duration of a single regulation block ranged from 9 to 45 s (mean = 28.68). Due to the hemodynamic nature of the BOLD signal, the duration of each regulation block in most of the studies (20 studies) ranged from 20 to 45 s and only 2 studies had shorter regulation blocks of 9 s and 12.5 s respectively (Veit et al., 2012; Zilverstand et al., 2015).

#### 3.5.2. Practice and transfer sessions

While a practice session is set to help participants familiarize the training procedure before the formal NF training, a transfer session is applied to examine maintenance of training effects on the neural and behavioral level in the absence of feedback. In the selected studies, only 5 included practice sessions (Berman et al., 2013; Rance et al., 2014a, 2014b; Tinaz et al., 2018; Zilverstand et al., 2015) and 4 of them reported successful NF training effects (Rance et al., 2014a, 2014b; Tinaz et al., 2018; Zilverstand et al., 2015). Eight studies included transfer sessions (Berman et al., 2013; Buyukturkoglu et al., 2015; Caria et al., 2007; Kanel et al., 2019; Rana et al., 2020; Ruiz et al., 2013; Tinaz et al., 2018; Yao et al., 2016) and 6 of them tested maintenance effects in a transfer session immediately following NF training (**Table 2**). More specifically, there was one study in Parkinson’s disease showing significant maintenance effects on self-regulation of the insula-dmFC functional connectivity in a training protocol lasting around 3 weeks (Tinaz et al., 2018) and 2 studies reporting significant maintenance effects on self-regulation of the insula activity in the transfer session immediately following NF training (Kanel et al., 2019) or 2 days after NF training (Yao et al., 2016). The latter study also reported that changes in rsFC of the AI but not in the behavioral performance induced by NF training were also maintained in the transfer run (Yao et al., 2016).

#### 3.5.3. Control group

Inclusion of a control group is necessary for excluding the possibility that the NF training effect is not specific for feedback of activity changes in target brain regions. However, in the 22 qualified studies, more than half (12 studies) did not include a control group. In the remaining 10 studies, sham feedback from an unspecific or task-irrelevant region is the most common manipulation (5 studies) used in the control group (Kanel et al., 2019; Karch et al., 2019, 2015; Lawrence et al., 2014; Yao et al., 2016), followed by 3 studies with no feedback (Linden et al., 2012; Tinaz et al., 2018; Zilverstand et al., 2015) and 2 studies using 2 control groups with sham or no feedback (Caria et al., 2010, 2007). Among the other 12 studies without a control group, there were 2 studies comprising 2 NF training groups comparing training difference between two different target ROIs (AI vs. ACC; Emmert et al., 2014) or of ability in self-regulation of the insula between lean and obese individuals (Frank et al., 2012). Interestingly, while 6 out of the 10 studies with control groups were published later than 2015, only 3 among the 12 studies without control groups were published later than 2015 (all details see **Table 2**), suggesting increasing methodological rigor in the field.

#### 3.5.4. Stimuli

In this review, 18 studies applied stimuli to participants during training, with most of the studies (14 studies) presenting participants with different types of visual stimuli, including emotional faces (Cohen Kadosh et al., 2016; Ruiz et al., 2013), emotional/threat-related stimuli from the IAPS (Caria et al., 2010; Johnston et al., 2010; Lawrence et al., 2014; Linden et al., 2012; Sitaram et al., 2014; Veit et al., 2012), painful pictures (Yao et al., 2016), symptom-related aversive scenes (Buyukturkoglu et al., 2015; Zilverstand et al., 2015), and addictive pictures (Karch et al., 2019, 2015; Rana et al., 2020). Another 3 studies used painful heat stimulation to induce activity of the insula during regulation (Emmert et al., 2014; Rance et al., 2014a, 2014b) and one study presented emotional auditory stimuli to aid implementation of regulation strategies by the participants (Kanel et al., 2019). These stimuli were simultaneously presented during regulation blocks to induce insula activation in the participants (Buyukturkoglu et al., 2015; Caria et al., 2010; Emmert et al., 2014; Kanel et al., 2019; Karch et al., 2019, 2015; Rana et al., 2020; Rance et al., 2014a, 2014b; Veit et al., 2012; Zilverstand et al., 2015), or were presented following regulation/rest blocks (Yao et al., 2016) or pre- and post-NF training (Buyukturkoglu et al., 2015; Lawrence et al., 2014; Ruiz et al., 2013; Sitaram et al., 2014) to measure training-induced behavioral changes. The remaining 4 studies did not use any stimuli (Berman et al., 2013; Caria et al., 2007; Frank et al., 2012; Tinaz et al., 2018). Details of stimuli presentation time points and stimuli types are presented in **Table 2**.

### 3.6. Outcome measurements

#### 3.6.1. NF training success on insula regulation

Training success on insula regulation was determined by comparing the difference in insula activity between the NF training and control groups, between the first and the last training sessions, or between regulation and rest blocks. Successful regulation of insula activity was reported in 19 of the 22 qualified studies (86.36%) based on these different comparison methods. The most consistent finding on training success was a significant difference in insula activity between the first and the last training sessions in the NF training group, which was found in 16 studies (including 10 studies and 6 studies in healthy and clinical populations separately; see **Fig. 2A**). Ten of the studies (some studies reported training success based on more than one comparison) reported significant differences between the regulation and rest blocks in the NF training group and 2 studies reported significant differences between the experimental and control groups (details see **Table 3**). Three studies did not show any significant training effects on insula regulation (Berman et al., 2013; Buyukturkoglu et al., 2015; Sitaram et al., 2014).

**Table 3.** Statistics of training success contrasts on insula self-regulation and associated functional connectivity changes.

| Study | Up/Down | EG vs. CG |  | First vs. Transfer | First vs. Last |  | Regulate vs. Rest |  | Functional connectivity |  |
| --- | --- | --- | --- | --- | --- | --- | --- | --- | --- | --- |
|  |  | Across sessions | Transfer | EG | EG | CG | EG | CG | Region | Findings |
| Berman et al. (2013) | Up | / | / | NS | NS | / | / | / | dmPFC | ↑ p < 0.0001 |
| Buyukturkoglu et al. (2015) | Down | / | / | NS | NS | / | - | / | / | / |
| Caria et al. (2007) | Up | - | - | NS | p < 0.012 | NS | p = 0.001 | / | / | / |
| Caria et al. (2010) | Up | NS | / | / | p = 0.012* | MI (p = 0.034) | / | / | / | / |
| Emmert et al. (2014) | Down | / | / | / | p < 0.05 | / | / | / | vlPFC | ↑ p < 0.05 |
| Frank et al. (2012) | Up | / | / | / | p < 0.05 | / | / | / | MCC/MTC | ↑ p < 0.001 |
| Johnston et al. (2010) | Up | / | / | / | p < 0.029 | / | p = 0.002 | / | / | / |
| Kadosh et al. (2016) | Up/Down | / | / | / | NS | / | p = 0.01 | / | Amygdala# | p = 0.001 § |
| Kanel et al. (2019) | Up | NS | p = 0.018 | NS | p = 0.041 | NS | / | / | / | / |
| Karch et al (2015) | Down | / | / | / | p = 0.048 | NS | / | / | mPFC/dlPFC | ↑ p < 0.05 |
| Karch et al (2019) | Down | / | / | / | / | / | p < 0 .05 | / | / | / |
| Lawrence et al. (2014) | Up | - | / | / | p = 0.03* | NS | / | / | / | / |
| Linden et al. (2012) | Up | / | / | / | p = 0.04 | / | / | / | / | / |
| Rana et al. (2020) | Down | / | / | - | p < 0.05 | / | / | / | / | / |
| Rance et al. (2014a) | Up/Down | / | / | / | p < 0.05 | / | p < 0.05 | / | / | / |
| Rance et al. (2014b) | Up/Down | / | / | / | p < 0.05 | / | p < 0.05 | / | / | / |
| Ruiz et al. (2013) | Up | / | / | NS | p = 0.02 | / | p < 0.05 | / | mPFC/Amygdala# | p < 0.01 |
| Sitaram et al. (2014) | Up | / | / | / | NS | / | / | / | PCC# | p < 0.05 |
| Tinaz et al. (2018) | Up | / | - | p = 0.009 | p < 0.05 | / | / | / | dmFC | ↑ p = 0.009 |
| Veit et al. (2012) | Up/Down | / | / | / | NS | / | p < 0.001 | / | dmPFC/vlPFC | ↑ p < 0.001 § |
| Yao et al. (2016) | Up | p = 0.041 | p = 0.006 |  | p = 0.011 | NS | / | / | dlPFC/dmPFC/IPL | ↑ p < 0.001 |
| Zilverstand et al. (2015) | Down | p < 0.05 | / | / | p < 0.05 | NS | p < 0.05 | NS | / | / |

*Note.* Abbreviations in alphabetical order: Transfer = no feedback; - = Did not report; \* = Significant interaction effects; # = Granger Causality Modeling; § = Significant effects of functional connectivity was found in up-regulation.

#### 3.6.2. Behavioral outcomes associated with training success

Behavioral effects associated with NF training were measured in 18 studies included in the present review using different types of behavioral or symptom-related questionnaires/ratings/tasks (**Table 4**). Similar to brain outcomes, significant behavioral effects were also determined by comparing the difference between the NF training and control groups, between pre- and the post-training, or between regulation and rest blocks in the NF training relative to the control groups.

**Table 4.**
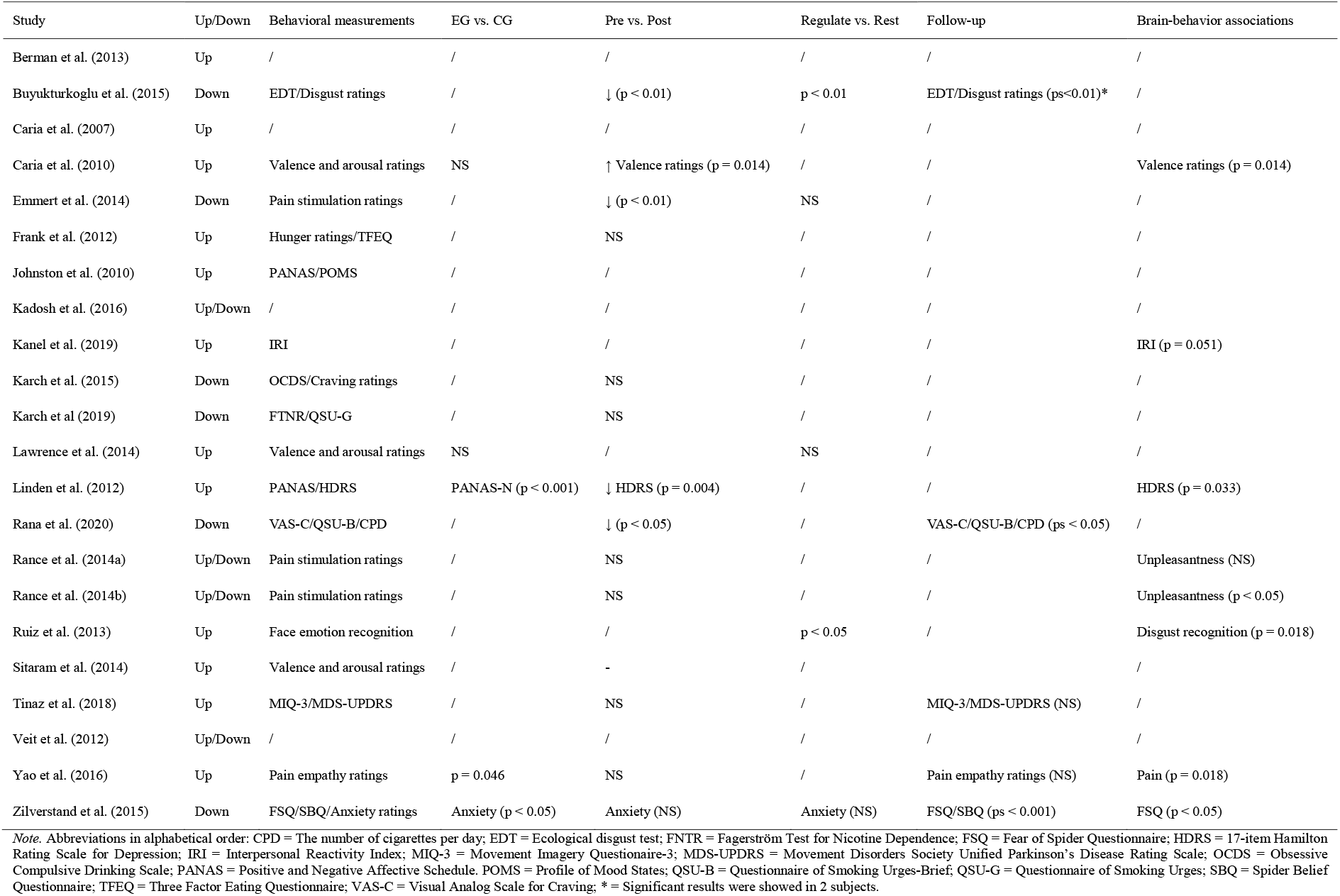
Statistics of behavioral measurement contrasts associated with NF training and brain-behavior associations.

| Study | Up/Down | Behavioral measurements | EG vs. CG | Pre vs. Post | Regulate vs. Rest | Follow-up | Brain-behavior associations |
| --- | --- | --- | --- | --- | --- | --- | --- |
| Berman et al. (2013) | Up | / | / | / | / | / | / |
| Buyukturkoglu et al. (2015) | Down | EDT/Disgust ratings | / | ↓ (p < 0.01) | p < 0.01 | EDT/Disgust ratings (ps<0.01)* | / |
| Caria et al. (2007) | Up | / | / | / | / | / | / |
| Caria et al. (2010) | Up | Valence and arousal ratings | NS | ↑ Valence ratings (p = 0.014) | / | / | Valence ratings (p = 0.014) |
| Emmert et al. (2014) | Down | Pain stimulation ratings | / | ↓ (p < 0.01) | NS | / | / |
| Frank et al. (2012) | Up | Hunger ratings/TFEQ | / | NS | / | / | / |
| Johnston et al. (2010) | Up | PANAS/POMS | / | / | / | / | / |
| Kadosh et al. (2016) | Up/Down | / | / | / | / | / | / |
| Kanel et al. (2019) | Up | IRI | / | / | / | / | IRI (p = 0.051) |
| Karch et al. (2015) | Down | OCDS/Craving ratings | / | NS | / | / | / |
| Karch et al (2019) | Down | FTNR/QSU-G | / | NS | / | / | / |
| Lawrence et al. (2014) | Up | Valence and arousal ratings | NS | / | NS | / | / |
| Linden et al. (2012) | Up | PANAS/HDRS | PANAS-N (p < 0.001) | ↓ HDRS (p = 0.004) | / | / | HDRS (p = 0.033) |
| Rana et al. (2020) | Down | VAS-C/QSU-B/CPD | / | ↓ (p < 0.05) | / | VAS-C/QSU-B/CPD (ps < 0.05) | / |
| Rance et al. (2014a) | Up/Down | Pain stimulation ratings | / | NS | / | / | Unpleasantness (NS) |
| Rance et al. (2014b) | Up/Down | Pain stimulation ratings | / | NS | / | / | Unpleasantness (p < 0.05) |
| Ruiz et al. (2013) | Up | Face emotion recognition | / | / | p < 0.05 | / | Disgust recognition (p = 0.018) |
| Sitaram et al. (2014) | Up | Valence and arousal ratings | / | - | / |  | / |
| Tinaz et al. (2018) | Up | MIQ-3/MDS-UPDRS | / | NS | / | MIQ-3/MDS-UPDRS (NS) | / |
| Veit et al. (2012) | Up/Down | / | / | / | / | / | / |
| Yao et al. (2016) | Up | Pain empathy ratings | p = 0.046 | NS | / | Pain empathy ratings (NS) | Pain (p = 0.018) |
| Zilverstand et al. (2015) | Down | FSQ/SBQ/Anxiety ratings | Anxiety (p < 0.05) | Anxiety (NS) | Anxiety (NS) | FSQ/SBQ (ps < 0.001) | FSQ (p < 0.05) |

*Note.* Abbreviations in alphabetical order: CPD = The number of cigarettes per day; EDT = Ecological disgust test; FNTR = Fagerström Test for Nicotine Dependence; FSQ = Fear of Spider Questionnaire; HDRS = 17-item Hamilton Rating Scale for Depression; IRI = Interpersonal Reactivity Index; MIQ-3 = Movement Imagery Questionnaire-3; MDS-UPDRS = Movement Disorders Society Unified Parkinson’s Disease Rating Scale; OCDS = Obsessive Compulsive Drinking Scale; PANAS = Positive and Negative Affective Schedule. POMS = Profile of Mood States; QSU-B = Questionnaire of Smoking Urges-Brief; QSU-G = Questionnaire of Smoking Urges; SBQ = Spider Belief Questionnaire; TFEQ = Three Factor Eating Questionnaire; VAS-C = Visual Analog Scale for Craving; \* = Significant results were showed in 2 subjects.

In the 18 studies, 8 found significant behavioral effects, including 3 studies in healthy and 5 studies in clinical populations. In the 3 healthy population studies, while (Yao et al., 2016) on up-regulation of AI found significantly higher pain empathy ratings in the NF training compared with the control groups, the other 2 studies found significantly increased valence ratings to aversive pictures in the last compared with the first training session during up-regulation (Caria et al., 2010) but decreased pain perception ratings to heat stimulation in the last training session relative to the functional localizer session during down-regulation (Emmert et al., 2014). In the 3 clinical studies on insula down-regulation, 2 studies reported decreased disgust levels to symptom-evoking stimuli in OCD (Buyukturkoglu et al., 2015) or decreased craving scores in addicted smokers in the post-test relative to pre-test (Rana et al., 2020) and one study found lower anxiety levels to spider pictures in individuals with spider phobia in the NF training relative to control groups (Zilverstand et al., 2015). In the remaining 2 studies on up-regulation, NF training induced symptom relief in depression as measured by mood and symptom severity questionnaires pre- and post-test (Linden et al., 2012) and improvement in recognition accuracy to disgust faces but impaired recognition accuracy to happy faces in schizophrenia patients in the regulation relative to the baseline blocks (Ruiz et al., 2013). All details of behavioral outcome measurements and findings are showed in **Table 4**.

In the 18 studies, 5 studies included behavioral follow-up measurements varying from 2 days to 6 months after NF training (Buyukturkoglu et al., 2015; Rana et al., 2020; Tinaz et al., 2018; Yao et al., 2016; Zilverstand et al., 2015) and 3 clinical studies found sustained decrease in disgust ratings to symptom-evoking stimuli in OCD (Buyukturkoglu et al., 2015), craving scores in addicted smokers (Rana et al., 2020) and symptom levels in individuals with spider phobia (Zilverstand et al., 2015) (**Table 4**).

#### 3.6.3. Brain outcomes associated with training success

In addition to training success on self-regulation of insula activity, 9 studies examined functional connectivity changes during NF training and only increased functional connectivity was found in all of these studies. Increased functional connectivity of the insula was found mainly in frontal regions and additionally in the cingulate cortex and amygdala (**Table 3**). Another study in which only one out of 4 subjects finished all of the NF training sessions, showed casual functional connectivity changes mainly driven by the posterior cingulate cortex (**Table 3**) and thus findings in this respect should be interpreted with caution given the small sample size (Sitaram et al., 2014).

For rsFC changes, there were only 2 studies reporting significant rsFC changes induced by NF training, with one study in healthy subjects reporting increased rsFC of the insula mainly with regions of the empathic network/mirror neuron system (MNS) but decreased rsFC with regions of the default mode network (DMN) (Yao et al., 2016) and the other in patients with alcohol use disorder showing increased rsFC of the insula mainly with frontal regions (Karch et al., 2015).

#### 3.6.4. Brain-behavior associations

Brain-behavior associations were reported in 8 studies involving both the healthy and clinical populations. There were 4 studies in healthy subjects demonstrating significant associations between NF-induced changes of the insula activity and different behavioral measurements including changes of valence ratings to aversive pictures (Caria et al., 2010), empathic ratings to painful pictures (Yao et al., 2016), unpleasantness ratings to painful heat stimulation (Rance et al., 2014b), and dispositional empathy levels as measured by the Interpersonal Reactivity Index (Kanel et al., 2019). In another 3 clinical studies, NF-induced insula activity was found to be significantly correlated with improvement in recognition accuracy of disgust faces (Ruiz et al., 2013), fear/belief of spiders as measured by the Fear of Spider Questionnaire and Spider Belief Questionnaire (Zilverstand et al., 2015), and depressive symptom severity as measured by the Hamilton Rating Scale for Depression and mood as measured by the Positive and Negative Affect Schedule (Linden et al., 2012) (**Table 4**).

## 4. Discussion

In the present systematic review, we obtained 22 qualified studies on insula regulation based on rtfMRI NF training. Participants’ demographics, feedback types, regulation strategies, localizing approaches, protocol design, outcome measurements and main findings were systematically reviewed to provide a comprehensive overview of progress in the field of insula rtfMRI NF training. We have organized the discussion of these 22 qualified studies in accordance with the three main objectives of this review.

### 4.1. What characteristics were used in NF training protocols among studies?

#### 4.1.1. Clinical vs. healthy populations

RtfMRI NF training targeting the insula has been applied in both healthy individuals and various clinical populations with mental disorders including MDD, OCD, addiction, spider phobia, as well as Parkinson’s disease (see **Table 1**). Based on the reviewed publications, the rate of training success on insula self-regulation was high in both healthy (92.31%) and the clinical populations (77.78%). Although the training success rate could be inflated by publication bias, the high training success rates can still indicate that the rtfMRI-based insula self-regulation is feasible in healthy subjects as well as clinical populations despite strong (meta)cognitive/affective deficits in the disorders included in the present study. This notion is also further corroborated by a recent systematic review reporting a training success rate of 89% of different target regions in 66 clinical studies (Tursic et al., 2020) and a meta-analysis study showing a medium-sized effect of rtfMRI NF training success in psychiatric disorders based on 17 studies (Dudek and Dodell-Feder, 2021). In another recent review study, training success rate was even higher in clinical compared to healthy populations (Haugg et al., 2021), reflecting a high dependence of calculated training success rate on sampling of publications and thus suggesting caution in interpreting those findings. Furthermore, the 22 qualified studies reporting training success encompassed individuals from a broad age range varying from 11.6 to 64.49 years, indicating that rtfMRI training on the insula is feasible for individuals across a broad age range.

#### 4.1.2. Continuous vs. intermittent feedback

The majority of the studies (18 studies) on insula regulation applied continuous feedback and only 4 studies used intermittent feedback (Buyukturkoglu et al., 2015; Rana et al., 2020; Tinaz et al., 2018; Zilverstand et al., 2015). Both types of feedback were effective (training success rate: 88.89% and 75% for continuous and intermittent feedback separately) in aiding subjects to gain the ability to regulate their insula activity. Although there are no studies having explored the superiority of the two types of feedback on insula regulation, intermittent feedback has been found to be superior in regulating the premotor cortex and amygdala as compared to continuous feedback in healthy subjects (Hellrung et al., 2018; Johnson et al., 2012). Another study in 14 tinnitus patients revealed that continuous feedback is more suitable for long-term NF training with multiple sessions and intermittent feedback is more efficient for single session NF training with respect to voluntary control over the auditory cortex (Emmert et al., 2017b). However, given the different numbers of studies between the two types of feedback in the present review and the small sample size (8-16 subjects in one single training/control group) in the previous studies directly comparing different feedback types the superiority of training efficacy between feedback types should be inferred with caution.

#### 4.1.3. Regulation strategies: specific instructions vs. free regulation

Selection of appropriate regulation strategies is critical for efficient regulation of brain activity during NF training. Among the 22 qualified studies, more studies preferred applying specific regulation strategies (15 studies) than free strategies (7 studies), although both of them were demonstrated to be efficient in regulating insula activity with a training success rate of 86.67% and 85.71% for specific and free regulation strategies, respectively. On the one hand, the advantage of providing a specific strategy is that it can provide a specific direction for subjects to work on and consequently may reduce trial and error periods at the beginning of training sessions, whereas it may have disadvantages where the provided strategy is invalid or not suitable for the individual or the specific population (e.g., Sepulveda et al., 2016). On the other hand, free strategies allow subjects to discover their own strategy for the specific target region, consequently leading to the most efficient strategy, however this approach comes at the cost that subjects may fail to find an efficient strategy. Furthermore, a 7T fMRI study has shown that different cognitive strategies used for pain attenuation can induce different patterns of brain activity in brain regions including the insula (Schulz et al., 2019). Thus, in the context of decreasing inter-subject variability induced by the use of different regulation strategies, a specific strategy, whose efficacy should be validated before application, may be more suitable for future studies, although it should be emphasized that currently there is no empirical evidence to confirm its superiority.

For regulation direction, we found ample evidence for both successful up- and down regulation of the insula activity, with a training success rate of 87.50% in 16 studies involving up-regulation and 70% in 10 studies involving down-regulation. In sharp contrast to up-regulation, most of the down-regulation studies (9 in 10 studies in down-regulation vs. 5 in 16 studies in up-regulation) presented stimuli during active regulation blocks to achieve a strong baseline of insula activity for down-regulation. Furthermore, free regulation strategies were used in most of the down-regulation as compared to up-regulation studies (7 in 10 studies in down-regulation vs. 4 in 16 studies in up-regulation), suggesting that determination of a specific regulation strategy may be more difficult for down-regulation studies. These findings are of informative and referential value for training protocol design in future studies.

#### 4.1.4. Protocol design

Participants were mostly successful in acquiring neural self-regulation in brief NF trainings, although training sessions can last from several sessions to days (Sitaram et al., 2017; Thibault et al., 2018). In the present review, training protocols ranged from 3 to 24 sessions, with clinical studies incorporating slightly more sessions relative to healthy studies (9.78 vs. 8 sessions). Clinical studies are normally designed to determine the therapeutic potential of rtfMRI NF training with respect to improving clinical symptoms. Separating multiple sessions into different days and repeating the individual sessions over days can thus help to avoid fatigue of patients and reinforce the training effect. In addition to active training sessions, some previous studies additionally included a practice/pre-training no-feedback session for participants to familiarize themselves with the training procedure and regulation strategies and this has been identified as a key factor in influencing subsequent training effects (Haugg et al., 2021). It has also been suggested that training success can be more reliable when a practice session is provided for participants to practice the instructed regulation strategy (Barreiros et al., 2019). A recent functional near-infrared spectroscopy NF study employing a practice session has also reported successful regulation of lateral orbitofrontal activity (K. Li et al., 2019). In the present review, there were 5 studies incorporating a practice session and 4 of them reported significant training effects. More emphasis should be paid on the benefits of using a practice/pre-training session in future studies.

In contrast to other factors, the control group does not affect training effects per se but is crucial to determine whether the training effect is NF-specific. In this review, less than half of the studies (10 studies) included a control group in the training protocol. Note that inclusion of a control group has become increasingly more applied in more recent studies, with 6 out of 9 studies published since 2015 but in contrast 4 out of 13 studies published before 2015 including a control group, although there were indeed some early studies encompassing a well-controlled group in their protocol design (e.g., Caria et al., 2010, 2007). It has been recommended that selection of a control group should be based on the specific goal of a NF training study (Sorger et al., 2019). Although it is ideal to control as many confounding effects as possible by using more control conditions, it is costly and sometimes not feasible in practice. The control groups used in the 10 studies varied from sham feedback from an unspecific or control region (Kanel et al., 2019; Karch et al., 2019, 2015; Lawrence et al., 2014; Yao et al., 2016), to no feedback (Linden et al., 2012; Tinaz et al., 2018; Zilverstand et al., 2015), or to both of the sham and no feedback controls (Caria et al., 2010, 2007), simultaneously being matched with instructed or free regulation strategies identical to the training group. A control group with matched regulation strategies is a more common method for control conditions that not only can minimize the number of control groups and maximize statistic power but also can simultaneously control for both placebo/unspecific effects and the use of regulation strategies (cf. Sorger et al., 2019).

In the present review, different types of stimuli were additionally presented in 18 studies and mostly visual including emotional/threat-related stimuli (Caria et al., 2010; Johnston et al., 2010; Lawrence et al., 2014; Linden et al., 2012; Sitaram et al., 2014; Veit et al., 2012), symptom-related/evoked scenes (Buyukturkoglu et al., 2015; Zilverstand et al., 2015), painful pictures/stimulation (Yao et al., 2016), and emotional auditory stimuli (Kanel et al., 2019). These stimuli are all within the scope of functional relevance of the insula and were presented in localization tasks to localize the target region, during regulation blocks to induce reactivity of the insula, or following regulation/rest blocks or pre- and post-NF training to measure training-induced behavioral changes based on different research goals in different studies (see **Table 2**). When and where to present stimuli in the training protocol should be determined based on research purpose and should avoid possible adverse effects such as distraction from performing regulation strategies.

### 4.2. Can rtfMRI NF training lead to consistently successful volitional control of insula, and if so, can this training effect be maintained to a “transfer” session without any neurofeedback?

Training success on insula self-regulation in the 22 qualified studies was determined by comparing difference of the insula activity between the NF training and control groups, between the first and the last training sessions, or between regulation and rest blocks in the NF training relative to the control groups. Based on these comparison methods, 19 of them (86.36%) reported successful self-regulation of insula, indicating that rtfMRI-based NF training is an efficient approach in aiding participants to acquire voluntary control over the insula activity. The most consistent method was comparison of the insula activity between the first and the last training sessions in the NF training group (16 of the 19 studies), reflecting improvement of self-regulation ability with progress over training sessions. Although there are very few studies having specifically reviewed publications of rtfMRI NF training on a specific single brain region or pathway, comparable training efficacy has been reported in recent review/meta-analysis studies on different target regions in both healthy and clinical populations (Dudek and Dodell-Feder, 2021; Thibault et al., 2018; Tursic et al., 2020). However, it has to be noted that given the publication bias, the reported ratio is not an indicator of training success in the whole population of studies, but rather an index of successful insula regulation in the 22 included studies.

Moreover, there is some preliminary but promising evidence for the maintenance of self-regulation ability on insula activity when no feedback information is available (Dudek and Dodell-Feder, 2021; Thibault et al., 2018). In the 22 included studies, the maintenance of insula self-regulation ability has been examined in 8 studies, with 2 studies reporting significant maintenance effects on insula regulation either in a transfer session immediately following NF training (Kanel et al., 2019) or 2 days after NF training (Yao et al., 2016). Another study focusing on functional connectivity-based NF training on the insula-dmFC pathway has also shown significant maintenance effects in patients with Parkinson’s disease (Tinaz et al., 2018). However, there were very few studies having explored maintenance effects lasting for days or even longer (cf., Tinaz et al., 2018; Yao et al., 2016; Zhao et al., 2019) and most studies (6 of 8 studies) tested the maintenance effect in a transfer session immediately following NF training. The maintenance of self-regulation ability for a longer period is of tremendous significance by suggesting the possibility of independent self-regulation on brain activity without continuous dependence on the MRI scanner and thus is key for a promising potential in clinical translational settings. In the present review, different types of stimuli were additionally presented in 18 studies and mostly visual including emotional/threat-related stimuli (Caria et al., 2010; Johnston et al., 2010; Lawrence et al., 2014; Linden et al., 2012; Sitaram et al., 2014; Veit et al., 2012), symptom-related/evoked scenes (Buyukturkoglu et al., 2015; Zilverstand et al., 2015), painful pictures/stimulation (Yao et al., 2016), and emotional auditory stimuli (Kanel et al., 2019). These stimuli are all within the scope of functional relevance of the insula and were presented in localization tasks to localize the target region, during regulation blocks to induce reactivity of the insula, or following regulation/rest blocks or pre- and post-NF training to measure training-induced behavioral changes based on different research goals in different studies (see **Table 2**). When and where to present stimuli in the training protocol should be determined based on research purpose and should avoid possible adverse effects such as distraction from performing regulation strategies.

### 4.3. What neural and behavioral changes have been induced by rtfMRI NF training on insula regulation?

#### 4.3.1. Behavioral changes induced by NF training

Behavioral/symptom-related changes associated with successful self-regulation of regional brain activity are a prerequisite for the therapeutic potential of rtfMRI NF training. There is ample evidence for changes in a variety of human behaviors or clinical symptoms associated with NF training in both healthy and clinical populations (Dudek and Dodell-Feder, 2021; Sulzer et al., 2013; Tursic et al., 2020). For example, in a recent review the majority of studies with behavioral measurements (44 of 59 studies) reported significant changes of behavioral indices induced by NF training (Thibault et al., 2018). In the present review, 18 studies measured behavioral/symptom-related changes yet only 8 of them reported significant changes associated with NF training, including ratings to emotional stimuli or painful stimulation in healthy subjects (Caria et al., 2010; Emmert et al., 2014; Yao et al., 2016) and improvement in symptom-related indices in psychiatric disorders (Buyukturkoglu et al., 2015; Linden et al., 2012; Rana et al., 2020; Ruiz et al., 2013; Zilverstand et al., 2015). Significant behavioral changes were observed following successful insula self-regulation in 7 of the 8 studies, except one study reporting decreased disgust ratings to aversive pictures in 2 of 3 OCD patients in the absence of successful down-regulation of insula activity (Buyukturkoglu et al., 2015). Not surprisingly, successful up- or down-regulation of insula activity corresponded with increase or decrease in behavioral/symptom-related measurements, respectively. Note that 10 of these 18 studies showed disassociations between successful self-regulation of the insula activity and training-induced behavioral changes, which could be derived from small sample sizes, difference of training protocol design or inadequate selection of behavioral indices.

Furthermore, behavioral changes were found to be correlated with NF-induced changes of the insula activity in 7 studies including both healthy (Caria et al., 2010; Kanel et al., 2019; Rance et al., 2014b; Yao et al., 2016) and clinical populations (Linden et al., 2012; Ruiz et al., 2013; Zilverstand et al., 2015). Significant brain-behavior associations provide more convincing evidence that behavioral/symptom-related changes are specifically associated with NF training-induced neural changes in the target region. Similar brain-behavior associations have also been found in previous rtfMRI studies on regulation of other brain regions such as the right inferior frontal cortex (Alegria et al., 2017; Rubia et al., 2019) and amygdala (Young et al., 2017b, 2017a) or of functional connectivity between the ventrolateral prefrontal cortex and amygdala (Zhao et al., 2019) and between the inferior frontal gyrus and inferior parietal lobe (Zweerings et al., 2019), suggesting a direct functional relevance of the neural modulation for those behavioral effects.

In addition, one study in healthy subjects (Yao et al., 2016) and 4 clinical studies (Berman et al., 2013; Rana et al., 2020; Tinaz et al., 2018; Zilverstand et al., 2015) conducted behavioral follow-up measurements varying from 2 days to 6 months after NF training. Three clinical studies among them revealed sustained decrease in symptom severity in patients with OCD (Buyukturkoglu et al., 2015), smoking addiction (Rana et al., 2020) and spider phobia (Zilverstand et al., 2015). These findings further verified the therapeutic potential of rtfMRI NF training in relieving symptoms in clinical populations characterized by insula dysfunction.

#### 4.3.2. Functional connectivity changes induced by NF training

In addition to changes of insula activity during NF training, 9 studies have also examined changes in functional connectivity of the insula and all revealed increased functional connectivity of the AI mainly with medial and dorsal lateral prefrontal regions and the cingulate cortex within the regulatory control network. Acquisition of voluntary control over brain activity by NF training is particularly involved in the regulation learning process and consequently the underlying regulatory and learning networks (Emmert et al., 2016). Similar increased functional connectivity with these networks has also been found in previous rtfMRI studies targeting other regions such as the amygdala in healthy subjects (Zotev et al., 2011), the orbitofrontal cortex in subjects with high contamination-related anxiety (Scheinost et al., 2013), and the auditory cortex in patients with chronic tinnitus (Haller et al., 2010). These findings suggest that successful regulation may share a similar neural modulation mechanism of functional coupling between different target regions and similar regulation and learning networks generally involved in NF control. Furthermore, functional connectivity between other target regions (e.g., amygdala and core language network nodes) and brain regions including the prefrontal cortex have been found to be correlated with NF training success (Young et al., 2018) and symptom improvement in patients with major depression and schizophrenia (Young et al., 2018; Zweerings et al., 2019). However, no studies to date have investigated whether functional connectivity changes of the insula during NF training associates with the aforementioned behavioral changes induced by training. Investigations on these associations can promote our understanding of how interaction between the target region and other brain regions during training may affect behavioral readouts or training efficacy beyond the NF-induced regional activity changes in isolated regions.

RsFC is another neural index measuring NF training effects on insula self-regulation, although the proportion of studies examining this index is rather low (2 out of 22 studies). In contrast to the more convergent findings on increased functional connectivity between the insula and the regulation network during NF training, preliminary findings in one study on rsFC changes induced by NF training showed increased rsFC mainly with regions of the empathic network/MNS and decreased rsFC with regions of the DMN in healthy subjects (Yao et al., 2016). In the other study, increased rsFC between the insula and frontal regions was found in patients with alcohol use disorder (Karch et al., 2015). Changes of rsFC induced by NF training have also been found in previous rtfMRI studies on other target regions (e.g., amygdala and the orbitofrontal cortex) either in healthy subjects (Li et al., 2016a) or subjects with MDD (Young et al., 2018; Yuan et al., 2014), PTSD (Misaki et al., 2018), and high contamination-related anxiety (Scheinost et al., 2013), indicating that intrinsic organization of brain regions can be altered by NF training and used as a reliable index for testing NF training effects.

## 5. Conclusion, limitations and future directions

In conclusion, the present review provides a systematic overview of progress in the field of insula regulation based on rtfMRI NF training by systemically reviewing participants’ demographics, feedback types, regulation strategies, localizing approaches, protocol design, outcome measurements and main findings in 22 qualified rtfMRI studies. The conclusions drawn based on the present overview need to be considered in the context of the following limitations: (a) heterogeneity of characteristics such as sample size, populations, feedback types, training protocols, regulation strategies across different studies did not allow a quantitative meta-analyses and thus a qualitative review was conducted; (b) the potential of a publication bias may not allow statistical inferences with respect to determining the most efficient training characteristics (see also Barreiros et al., 2019). Despite these limitations, our systematic review can still promote our understanding of mechanisms underlying rtfMRI-based insula regulation, its associated behavioral and neural changes, how to optimize selection of characteristics and principles of NF protocol design.

Based on findings of the present study as well as previous studies, we suggest a number of guidelines for future studies targeting the insula in particular as well as other regions in general. With respect to feedback types, a continuous feedback may be preferential for NF training comprising multiple sessions and long-term multiple training sessions should be divided into brief sessions when being applied in clinical patients to avoid fatigue and reinforce the positive training effects. Furthermore, in comparison to free regulation strategies, specific strategies that have been validated before application are recommended. Future studies can also consider including a practice session for subjects to become familiar with the training procedure and practice/validate the regulation strategy. For target regions with distinguishable landmarks such as the insula, functional localization is adequate, whereas for sub-cortical regions without a clear landmark, a combination of functional and anatomical approaches can achieve more accurate localization of the target ROI. In addition, although design of training protocols such as whether a transfer session and stimuli are used should be based on specific research purpose, training protocol design, statistical analyses and reporting of results following common criteria and a priori determination of sample and effect sizes as well as pre-registration of studies are highly recommended (e.g., Fede et al., 2020; Ros et al., 2020). Progress in rtfMRI research standardization will enable comparison and quantitative meta-analysis across different studies and thus be more informative in optimizing future NF training protocol design and determining key factors influencing training efficacy. More advanced rtfMRI NF techniques such as decoded NF based on Multi-Voxel Pattern Analysis and functional connectivity-based NF based on Dynamic Causal Modeling can also bring new advances in this field (Koush et al., 2013; Watanabe et al., 2017). Taken together, we believe our review will inspire and inform both fundamental research and therapeutic translation of rtfMRI NF training as an intervention in mental disorders particularly those with insula dysfunction.

## Acknowledgements

This work was supported by the National Key Research and Development Program of China (grant number: 2018YFA0701400) and the National Natural Science Foundation of China (NSFC) grants (grant number: 31700998).

